# Mapping the human corticoreticular pathway with multimodal delineation of the gigantocellular reticular nucleus and high-resolution diffusion tractography

**DOI:** 10.1101/2021.08.27.457996

**Authors:** Pierce Boyne, Mark DiFrancesco, Oluwole O. Awosika, Brady Williamson, Jennifer Vannest

## Abstract

The corticoreticular pathway (CRP) is a major motor tract that provides volitional input to the reticular formation motor nuclei and may be an important mediator of motor recovery after central nervous system damage. However, its cortical origins, trajectory and laterality are incompletely understood in humans. This study aimed to map the human CRP and generate an average CRP template in standard MRI space. Following recently established guidelines, we manually delineated the primary reticular formation motor nucleus (gigantocellular reticular nucleus [GRN]) using several group-mean MRI contrasts from the Human Connectome Project (HCP). CRP tractography was then performed with HCP diffusion-weighted MRI data (N=1,065) by selecting diffusion streamlines that reached both the frontal cortex and GRN. Corticospinal tract (CST) tractography was also performed for comparison. Results suggest that the human CRP has widespread origins, which overlap with the CST across most of the motor cortex and include additional exclusive inputs from the medial and anterior prefrontal cortices. The estimated CRP projected through the anterior and posterior limbs of the internal capsule before partially decussating in the midbrain tegmentum and converging bilaterally on the pontomedullary reticular formation. Thus, the CRP trajectory appears to partially overlap the CST, while being more distributed and anteromedial to the CST in the cerebrum before moving posterior to the CST in the brainstem. These findings have important implications for neurophysiologic testing, cortical stimulation and movement recovery after brain lesions. We expect that our GRN and tract maps will also facilitate future CRP research.

**HIGHLIGHTS:**

- The corticoreticular pathway (CRP) is a major tract with poorly known human anatomy
- We mapped the human CRP with diffusion tractography led by postmortem & animal data
- The CRP appears to originate from most of the motor cortices and further anterior
- The estimated CRP had distributed and bilateral projections to the brainstem
- These findings have important implications for motor recovery after brain lesions

## 1. INTRODUCTION

The cortico-reticulo-spinal tract (CReST) is the original motor pathway.[1] It has been providing the primary descending commands for fundamental upper and lower limb movement synergies (e.g. reaching) and central pattern generation (e.g. locomotion) since before the evolution of the corticospinal tract (CST) and other motor projections.[1] Across evolution, the CReST has been conserved and further developed,[2] remaining among the fastest conducting motor pathways even in primates.[1] The corticoreticular pathway (CRP) segment of the CReST projects from distributed regions of the cerebral cortex to provide volitional input to the reticulospinal tract (RST) nuclei in the medial pontomedullary reticular formation.[3–7] RST neurons project to spinal cord interneurons and lower motor neurons, forming both excitatory and inhibitory synapses that span multiple joints and limbs.[2]

Although it has received much less attention than the CST,[8,9] there is now burgeoning interest in the CReST as a potential mediator of commonly observed movement patterns[9–12] and motor recovery[13–26] after central nervous system damage. For example, the distributed projections of the CRP offer the possibility of a proximal bypass around brain lesions like stroke, to potentially compensate for disruption of motor pathways and restore some volitional movement function.[7,10,17–20] However, the CRP has been difficult to study in humans, partly because its anatomy has not been fully charted.

MRI-based diffusion tractography is a non-invasive method that estimates tract trajectories by simulating streamlines through pre-processed diffusion-weighted images.[27–30] One research group has begun mapping the healthy human CRP using this method (Table 1).[31–34] Primarily focusing on CRP origins in the secondary motor cortices (supplementary motor area & premotor cortex), studies from this group have reported a CRP trajectory through the superior corona radiata, posterior limb of the internal capsule (anteromedial to the CST) and midbrain tegmentum (posterior to the CST) to the area of the pontomedullary reticular formation.[31,34]

**Table 1.** Summary of prior studies assessing normal human corticoreticular pathway (CRP) anatomy compared with the current study.

| Reference | N | Diffusion MRI acquisition | Tractography method | Cortical streamline inclusion regions | Brainstem streamline inclusion regions | Estimated CRP origins and/or trajectory |
| --- | --- | --- | --- | --- | --- | --- |
| Yeo 2012[31] | 24 | 1.5T, 6-ch coil, 2.3 mm isotropic voxels, 32 diffusion directions at b=1000 s/mm<sup>2</sup> | FSL probtrackx software (ball and stick model) using individual data | Premotor*† | Medullary RF (seed),†‡ midbrain tegmentum†‡ (ipsilateral only) | Trajectory through superior corona radiata and PLIC (anterior to CST), midbrain/pontine tegmentum to medullary RF |
| Yeo 2014[32] | 75 | 1.5T, 8-ch coil, 2.3 mm isotropic voxels, 32 diffusion directions at b=1000 s/mm<sup>2</sup> | DTI-Studio software (tensor model) using individual data | None (unrestricted cortical search window) | Medullary RF (seed),†‡ midbrain tegmentum†‡ (ipsilateral only) | Not described and difficult to discern from limited images provided |
| Jang 2014[33] | 42 | 1.5T, 8-ch coil, 2.3 mm isotropic voxels, 32 diffusion directions at b=1000 s/mm<sup>2</sup> | FSL probtrackx software (ball and stick model) using individual data | S1†, M1†, premotor*†, & prefrontal area 8† (one gyrus anterior to premotor cortex) | Medullary RF (seed),†‡ midbrain tegmentum†‡ (ipsilateral only) | Origins found from all cortical regions tested. Streamline volumes from premotor & M1 were greater than S1 and prefrontal area 8. |
| Jang 2015[34] | 33 | 1.5T, 8-ch coil, 2.3 mm isotropic voxels, 32 diffusion directions at b=1000 s/mm<sup>2</sup> | FSL probtrackx software (ball and stick model) using individual data | Premotor*† | Medullary RF (seed),†‡ midbrain tegmentum†‡ (ipsilateral only) | Trajectory through superior corona radiata and PLIC (anteromedial to CST), midbrain/pontine tegmentum to medullary RF |
| HCP data[51] & methods for current study | 1,065 | 3T, 32-ch coil, 1.25 mm isotropic voxels, 90 diffusion directions each at b=1000, 2000 & 3,000 s/mm<sup>2</sup> | DSI Studio software (GQI & QSDR model-free method) using averaged data in MNI space | Frontal lobe (using automated labelling of gray/white matter cortical boundary surface) | GRN (labelled using contrast boundaries on multimodal high-resolution images; ipsilateral & contralateral) | Origins found from M1, SMA, preSMA, PMd, mPFC and aPFC. Trajectory through anterior & superior corona radiata, ALIC & PLIC (anteromedial to CST), midbrain/pontine tegmentum to medullary RF. Partial decussation in midbrain tegmentum. |

\*Premotor cortex label appears to cover SMA, preSMA, PMd and PMv. †Manually drawn on low resolution (2.3 mm isotropic acquisition) b0 image. ‡Contrast boundaries for region of interest not visible on b0 image and region location/size appears variable between references.
ALIC, anterior limb of internal capsule; aPFC, anterior prefrontal cortex; CST, corticospinal tract; GQI, generalized q-sampling imaging; HCP, human connectome project; M1, primary motor cortex; mPFC, medial prefrontal cortex; PLIC, posterior limb of internal capsule; PMd, dorsal premotor cortex; PMv, ventral premotor cortex; QSDR, Q-space diffeomorphic reconstruction; RF, reticular formation; S1, primary somatosensory cortex; SMA, supplementary motor area.

**Table 2.** Gigantocellular reticular nucleus (GRN) delineation details

| Table 2. Gigantocellular reticular nucleus (GRN) delineation details |  |  |  |
| --- | --- | --- | --- |
| GRN rostrocaudal zone | GRN boundary | Boundary structures | Primary image for delineation / boundary structure appearance |
| From superior aspect of GRN (in same axial plane as superior aspect of 7N; z = -34 mm in MNI coordinates) to inferior aspect of 6N (z = -35 mm) | Posterolateral | IRt (diagonal line between 6N & 7N) | T2w / hyperintense (6N, 7N) |
|  | Posterior | 6N, Pr, mlf | T2w / hyperintense (6N, Pr)<br>FA / hyperintense (mlf) |
|  | Medial | mlf, ts | FA / hyperintense |
|  | Anterior | ml, ctg | FA / hyperintense |
| From just inferior to 6N (z = -36 mm) to inferior aspect of 7N (z = -39 mm); GRN expanded laterally and posteriorly in this zone | Posterolateral | IRt (diagonal line between Pr & 7N) | T2w / hyperintense (Pr, 7N) |
|  | Posterior | Pr | T2w / hyperintense |
|  | Medial | mlf, ts | FA / hyperintense |
|  | Anterior | ml, ctg | FA / hyperintense |
| From just inferior to 7N (z = -40 mm) to inferior pons (z = -46 mm); GRN narrowed laterally and expanded anteriorly in this zone | Posterolateral | IRt (diagonal line between Pr & st) | T2w / hyperintense (Pr)<br>FA / hyperintense (st)<br>DEC / blue (st) |
|  | Posterior | Pr | T2w / hyperintense |
|  | Medial | mlf, ts | FA / hyperintense |
|  | Anterior | ml, ctg, IOC | FA / hyperintense (ml, ctg)<br>FA / hypointense (IOC) |
| From inferior aspect of pons (z = -47 mm) to inferior aspect of GRN (just superior to the rostral pole of Gr and where the central canal begins to form; z = -55 mm); GRN moved lateral as ml expanded and GRN posterior border moved anterior as 10N & 12N emerged in this zone | Posterolateral | IRt (diagonal line between 10N/12N & st) | T2w / hyperintense (10N, 12N)<br>FA / hyperintense (st)<br>DEC / blue (st) |
|  | Posterior | 10N/12N | T2w / hyperintense |
|  | Medial | mlf, ts | FA / hyperintense |
|  | Anterior | IOC | FA / hypointense |
| 6N, abducens nerve nucleus; 7N, facial nerve nucleus; 10N, vagus nerve nucleus; 12N, hypoglossal nerve nucleus; ctg, central tegmental tract; Gr, gracile nucleus; IOC, inferior olivary complex; IRt, intermediate reticular zone; mlf, medial longitudinal fasciculus; Pr, prepositus nucleus; st, spinothalamic tract; ts, tectospinal tract. |  |  |  |

Other researchers have begun using the methods and findings from these normative studies to inform CRP tractography among patients with brain pathology.^e.g.^[20–23,35] For example, most subsequent clinical studies have followed the prior method of mapping CRP projections exclusively from the secondary motor cortices,^e.g.^[22,23,35] and/or have iteratively redefined their tractography methods to obtain CRP maps similar to prior work.^e.g.^[21] Unfortunately, the previous estimates of the healthy human CRP that are informing clinical research have several important and unresolved discrepancies from more definitive animal studies using axonal tracing and neuronal recording.

First, invasive animal studies (including in primates) consistently report important CRP projections originating from the primary motor cortex.[3–7,14,19,36–39] These projections have also been observed in humans with diffusion tractography,[33] but are still often omitted from the search window in human CRP studies. In addition, a recent large-scale (N=607) axonal tracing study in the mouse confirmed the presence of additional CRP projections originating far anterior to the motor cortices in the medial prefrontal and anterior cingulate regions.[7] These projections increase the probability of at least partial CRP sparing after a brain lesion and could generate novel targets for neuromodulation. Preliminary evidence also suggests the medial prefrontal and anterior cingulate cortices could be relevant to human motor function,[40–42] including in aging,[43] stroke[44] and Parkinson Disease.[45] However, no prior human studies have tested for CRP projections from the medial prefrontal or anterior cingulate cortices.

Another discrepancy is that prior human studies have only tested for ipsilateral CRP projections,[31–34] whereas invasive animal studies (including in primates) have consistently identified bilateral CRP projections.[5–7,36,46,47] After unilateral stroke in mice, upregulation of contralateral CRP projections from the contralesional cortex to the ipsilesional brainstem appears to be an important recovery mechanism.[17] Upregulation of the contralesional CRP has also been reported in humans after stroke,[11,20] but the existence of contralateral CRP projections has not been previously assessed in our species.

An additional challenge for human CRP tractography has been that the brainstem target (reticular formation motor nuclei) is not currently available in any standard-space MRI atlas, and is poorly defined on most MRI acquisitions,[48] especially the low-resolution b0 image used for manual labeling in prior CRP studies.[31–34] This has presumably forced researchers to guess at the target boundaries and/or to iteratively redefine them based on the tractography results. Thus, it is not surprising that the reticular formation image labels used for CRP tractography appear to vary widely across previous studies.[20–23,31,33,35]

The current study aimed to overcome these prior limitations and generate a more accurate map of the normal human CRP. Using recently established guidelines,[48] we manually traced the primary reticular formation motor nucleus in standard MRI space using group-mean high-resolution multimodal contrast maps from the Human Connectome Project (HCP; N=1,065-1,096 across maps),[49] with reference to a histologic atlas[50] and an ultra-high resolution post-mortem brainstem MRI annotation.[48] We then performed tractography using HCP high-resolution diffusion-weighted MRI data (N=1,065),[51] that had been preprocessed, reconstructed and averaged in standard MRI space.[30] We used the entire frontal lobe as the cortical target region to allow for the possibility of CRP streamlines originating from the medial prefrontal or anterior cingulate areas. We also used this same target region when mapping the CST for comparison. In addition, we tested for the possibility of contralateral CRP projections by including bilateral brainstem targets when mapping each CRP. Finally, we used the results to generate the first human average template of the CRP in standard MRI space, which will be made publicly available at https://balsa.wustl.edu/study/show/v83MM upon acceptance of this manuscript for publication.

## 2. MATERIAL AND METHODS

### 2.1. Diffusion-weighted MRI acquisitions

The brain MRI data used for this analysis were obtained from the Human Connectome Project (HCP) 1200-subject release, which is publicly available at https://db.humanconnectome.org/.[52] This included diffusion-weighted imaging from 1,065 participants (29 ± 4 years old [mean ± SD]; 54% female) scanned on a Siemens 3T Skyra scanner, using a 2D spin-echo single-shot multiband echo-planar imaging (EPI) sequence (multi-band factor 3; 1.25 mm isotropic voxel size; repetition time, 5500 ms; echo time, 89.50 ms). In addition to 6 b0 images, 90 diffusion sampling directions were acquired at each of three b-values (1000, 2000 and 3000 s/mm2).[51]

### 2.2. Diffusion MRI preprocessing

HCP preprocessing included correction for EPI distortion, eddy currents, head motion and gradient non-linearity[53] (for updated details see https://humanconnectome.org/study/hcp-young-adult/document/1200-subjects-data-release/). Yeh and colleagues then reconstructed the diffusion data from each participant in a standard Montreal Neurological Institute (MNI) space,[30] which was a cropped version of the ICBM152 2009c nonlinear asymmetric template (MNI152NLin2009cAsym) from https://www.bic.mni.mcgill.ca/ServicesAtlases/ICBM152NLin2009. This was done using generalized q-sampling imaging and q-space diffeomorphic reconstruction,[54] which combines nonlinear spatial registration and high-angular-resolution reconstruction of diffusion data while preserving the directional information and the continuity of fiber geometry. Such preservation enables the use of tractography on group average data for an improved signal to noise ratio.[54] The outputs of this reconstruction were MNI-registered maps of spin distribution functions (SDF) at each voxel, for each participant, with 1 mm^3^ resolution. The SDF includes a spherical isotropic component and protruding peaks for each prominent diffusion direction within the voxel. Quantitative anisotropy (QA) values quantify the amount of anisotropic spins diffusing along each direction relative to the isotropic background diffusion.[55] SDFs were scaled so that free water SDFs normalized to one.[54] Each voxel was then averaged across participants to create a group-average SDF map,[30] which is publicly available at http://brain.labsolver.org/diffusion-mri-templates/hcp-842-hcp-1021 (HCP-1065 1-mm template).

### 2.3. Delineating tractography regions of interest

#### 2.3.1. Gigantocellular reticular nucleus (GRN) target

The brainstem target for CRP tractography was manually created to cover the GRN, since this is the primary nucleus that gives rise to the reticulospinal tracts.[2,56–58] To delineate the GRN, we viewed contrast boundaries on several high-resolution group-average images in MNI space that were generated from the same HCP 1200 subject release as the SDF map used for tractography. This included a fractional anisotropy (FA) image and a direction-encoded color (DEC) image from the primary diffusion orientation (both 1.0 mm isotropic; 1,065-participant averages; distributed within the FMRIB Software Library (FSL)[59]; available at: https://fsl.fmrib.ox.ac.uk/fsl/fslwiki/FSL). It also included structural T1w, T2w and T1w/T2w images (all 0.7 mm isotropic; 1,096-participant averages; available at: https://balsa.wustl.edu/gKm1).

The FA & DEC mean images included the exact same participants as the tractography SDF map. The structural mean images included 1,064 (99.9%) of the participants used to generate the tractography SDF map (one participant had diffusion-weighted images but not structural images), plus 32 additional participants who had structural but not diffusion-weighted images. The demographic distribution of the participants who contributed to the structural mean images was identical to that for the FA, DEC & SDF maps (29 ± 4 years old; 54% female).

The GRN delineation was guided by several sources, including: 1) the Paxinos brainstem atlas;[50] 2) recently published MRI segmentation rules for the GRN (https://fibratlas.univ-tours.fr/mediawiki/index.php/Gigantocellular_reticular_nucleus) and its boundary nuclei (http://fibratlas.univ-tours.fr/mediawiki/index.php);[48] and 3) a companion three-dimensional GRN annotation on ultra-high-resolution post-mortem brainstem images (T2w and FA) from 11.7T MRI (WIKIBrainStem; https://fibratlas.univ-tours.fr/?page_id=53).[48] We rigidly rotated and translated the WIKIBrainStem images to roughly match the MNI brain orientation, so that WIKIBrainStem slices could be referenced in a similar plane to the HCP group-average images during manual tracing.

As a starting point for the GRN delineation, we also nonlinearly registered the WIKIBrainStem GRN label to the HCP group-average images in MNI space. The nonlinear warp field was calculated from the WIKIBrainStem FA map to the HCP-1065 FA map using the FSL ‘FLIRT’[60] and ‘FNIRT’[61] algorithms, with a brainstem mask (dilated 2 mm) to only consider that region of the HCP FA map during registration. The calculated warp field was then used to align the WIKIBrainStem GRN label to the HCP images for further manual editing. There are slight differences between the MNI152NLin2009cAsym template (the tractography space) and the HCP MNI-space average images. Thus, we also calculated a nonlinear warp from the HCP average T1w image to the MNI152NLin2009cAsym T1w image (using FSL FLIRT and FNIRT), then applied the warp to the edited GRN label to align it more precisely to the tractography space.

#### 2.3.2. Corticospinal tract (CST) waypoint targets

To enable comparative CST tractography, we manually delineated two brainstem CST waypoints on the HCP group-average MNI-space images, primarily using the FA and DEC contrasts, with reference to the Paxinos brainstem atlas.[50] Each waypoint was drawn on four consecutive 1-mm axial slices. One was drawn at the level of the pons (z=−32 to −35 mm) and the other in the medulla (z=−49 to −52 mm). The nonlinear warp applied to the GRN label was then also applied to the CST waypoints, to more precisely align them with the MNI152NLin2009cAsym tractography space.

#### 2.3.3. Frontal cortex target

We included the entire frontal cortex as a tractography target because a recent axonal tracing analysis in the mouse found CRP projections from widespread frontal regions.[7] The parietal lobe was omitted to minimize false inclusion of ascending sensory fibers in the CRP & CST maps, and because both CRP & CST projections from the somatosensory cortex are relatively minor and similar.[7] To precisely delineate the frontal cortex of the MNI152NLin2009cAsym template brain, we first used FreeSurfer software v6.0[62] to automatically label regions of the cortex at their inner boundary with the white matter surface,[51,63] based on the Desikan-Killiany atlas (https://surfer.nmr.mgh.harvard.edu/fswiki/CorticalParcellation).[63,64] We then combined all such cortical surface labels belonging to the frontal lobe.

However, one of these labels is the paracentral lobule, which includes part of the somatosensory cortex in the parietal lobe. To remove this region from our frontal lobe map, we used the HCP Multimodal Parcellation v1.0[65] (available at: https://balsa.wustl.edu/976l8) to mask out Brodmann areas 1-3, 5 & 23. This operation required pre-registration of the MNI152NLin2009cAsym cortical surface mesh to the Conte69 population average meshes[66] used for the Multimodal Parcellation. Surface registration was done using Connectome Workbench v1.5.0 (https://www.humanconnectome.org/software/connectome-workbench) and the ‘MSM-Sulc’ multimodal surface matching algorithm.[67]

We then projected the frontal lobe gray/white matter boundary surface back into the template volume space and dilated it inwards by 2 mm into the white matter. This slight dilation was done to minimize false negatives from premature streamline termination due to the dense tangential fibers and sharp fiber turns present just below the cortex.[68–70] Finally, we ensured that the dilation did not encroach on any subcortical gray matter structures by dilating the relevant FreeSurfer volumetric segmentation labels[71] by 2 mm and masking them out of the frontal lobe map.

#### 2.3.4. Other tractography regions

We also generated several other regions using FreeSurfer volumetric segmentation labels[71] for the MNI152NLin2009cAsym template brain, to initiate or restrict tractography. This included maps of the cerebellum, corpus callosum, the cerebral cortex and a streamline seeding region comprised of the cerebral white matter and brainstem.

### 2.4. Diffusion tractography

Diffusion tractography was performed on the HCP-1065 group-average SDF map with DSI Studio software (http://dsi-studio.labsolver.org). This tractography algorithm leverages the QA values from the SDF to resolve multiple fiber populations within a voxel, filter out unlikely fiber directions and account for partial volume effects.[55] This method has been shown to outperform other diverse tractography methods by achieving the highest number of valid connections in an open competition.[72] For each tractography run, streamlines were seeded from every cerebral white matter and brainstem voxel, the propagation step size randomly varied from 0.5 to 1.5 mm[73] and the QA threshold randomly varied from 0.5 to 0.7 times Otsu’s threshold.[73] Streamlines were terminated upon entering the cerebral cortex to prevent false sulcal/gyral crossing.[74] Streamlines were discarded if they entered the cerebellum or corpus callosum, or if they were shorter than 40 mm or longer than 150 mm. Each tractography run continued until 50,000 successful streamlines were obtained.

#### 2.4.1. CRP-specific tractography methods

For the CRP, the angular threshold was 60 degrees and streamlines were considered successful if they reached both the frontal lobe and the GRN. The left and right CRPs were separated by selecting streamlines touching either the left or right frontal lobes respectively. This allowed for the possibility of tracing bilateral CRP projections (e.g. from the right frontal lobe to the left and right GRN), as consistently observed in animal neuronal tracing studies (including non-human primates).[5–7,36,46,47]

#### 2.4.2. CST-specific tractography methods

For the CST, we used a lower angular threshold of 45 degrees to avoid projections from the ventral premotor cortex. CST streamlines were considered successful if they reached the frontal lobe and both CST waypoints (in the pons and medulla). The left and right CSTs were separated by selecting streamlines touching all of the left or right regions of interest, to match the known CST trajectory.[8,75] For example, the right CST streamlines had to reach the right frontal lobe, right pontine CST waypoint and right medullary CST waypoint, which was superior to the pyramidal decussation.

### 2.5. Tractography post-processing

We generated volumetric streamline density images for each tract with DSI Studio. These images contain the number of successful streamlines in each brain voxel. Given the common use of the MNI152 nonlinear 6^th^ generation template distributed within FSL (e.g. for MNI-space registration of HCP data[53]), we aligned the streamline density images to that template space. As before, this nonlinear registration warp was calculated from the MNI152NLin2009cAsym T1w image to the FSL MNI152 T1w image using FSL FLIRT and FNIRT.

For better visualization of cortical streamline endpoints, we also projected the streamline density data onto an inflated surface model of the cerebral cortex. This was done by dilating the streamline density images by 3 mm, mapping the resultant images onto the gray/white matter boundary surface, and viewing the results on an inflated version of the cortical surface, that had vertex correspondence with the gray/white matter boundary surface. To facilitate localization of the results, we also projected the volumetric labels for different motor cortex subregions of interest from Mayka et al[76] onto the same cortical surface model. This included the primary motor cortex (M1), supplementary motor area (SMA), preSMA, and dorsal premotor region (PMd). The HCP Multimodal Parcellation[65] was then used to refine the motor subregion surface labels and add labels for the medial prefrontal cortex (mPFC), anterior prefrontal cortex (aPFC) and dorsolateral prefrontal cortex (DLPFC).

## 3. RESULTS

### 3.1. Delineation of tractography regions of interest

Figure 1 shows the WIKIBrainStem GRN label[48] alongside our manual tracing of the GRN and CST waypoints on multimodal HCP group-average images. Table 1 details how each GRN border was delineated and how its shape changed along its rostrocaudal extent. GRN borders in the horizontal plane were all identifiable by contrast boundaries on at least one of the HCP images, except the posterolateral border with the intermediate reticular zone in some slices, which had to be inferred based on visible landmarks and its position in other slices. In cases of voxel uncertainty in this region, we erred towards a wider posterolateral GRN label, since the intermediate reticular zone also contains reticulospinal neurons, especially near its border with the GRN.[2,56,57]

**Figure 1.**
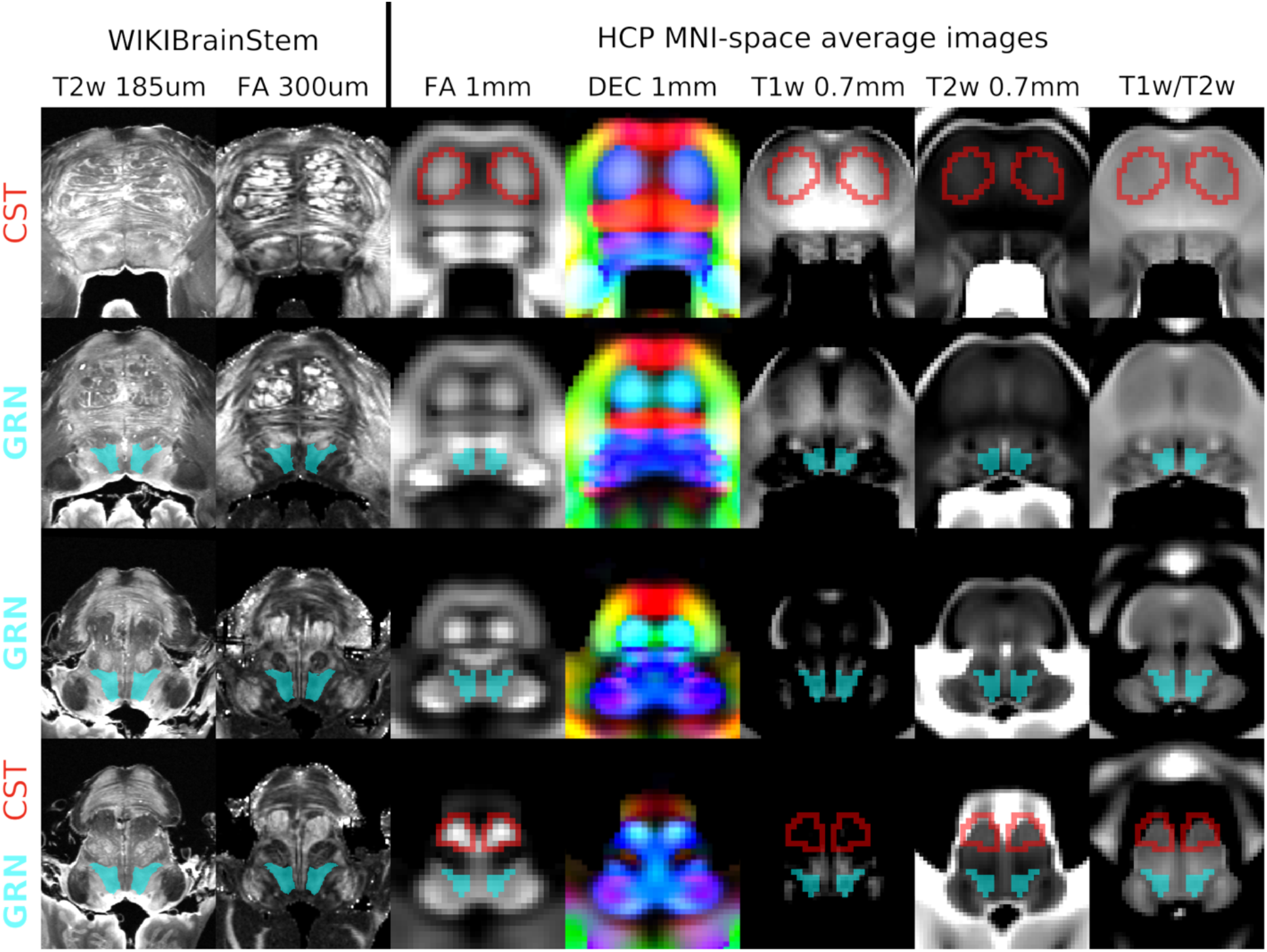
Delineation of the gigantocellular reticular nucleus (GRN) and corticospinal tract (CST) waypoints. The left two columns show the WIKIBrainStem ultra-high resolution 11.7T MRI contrasts and GRN annotation on a post-mortem brainstem,[48] after rigid rotation to the approximate orientation of the MNI template brainstem. The right columns show Human Connectome Project (HCP) high resolution 3T MRI contrasts averaged across 1,065 participants (FA, DEC) or 1,096 participants (T1w, T2w and T1w/T2w) after nonlinear alignment to the MNI template brain.[53] Each row shows approximately the same axial slice across all images (row 1, z = −32 mm in MNI space; row 2, z = −40 mm; row 3, z = −45 mm; row 4, z = −49 mm). Image intensity scaling was set to maximize the relevant contrast boundaries. We manually traced the GRN label (shown in translucent light blue) and the CST waypoints (outlined in red) on the HCP MNI-space average images. For clarity, labels are not shown on the DEC image. FA, fractional anisotropy; DEC, direction-encoded color. Data at: https://balsa.wustl.edu/G3xZg (upon acceptance for publication).

Likewise, the superior and inferior GRN boundaries were not well defined, and we erred towards a longer rostrocaudal extent for the GRN label, knowing that its superior neighbor (caudal pontine reticular nucleus) and inferior neighbor (ventral medullary reticular nucleus) both also contain reticulospinal neurons.[2,56,57] Conversely, some narrow isthmuses of the WIKIBrainStem GRN label were excluded from the HCP GRN delineation because they were narrower than the voxel width. This was in the pons around z = −34 to −38 mm in MNI coordinates.

The CST waypoint borders were all easily identifiable on the FA and DEC images. Figure 2 shows all of the tractography regions in the MNI152NLin2009cAsym tractography space.

**Figure 2.**
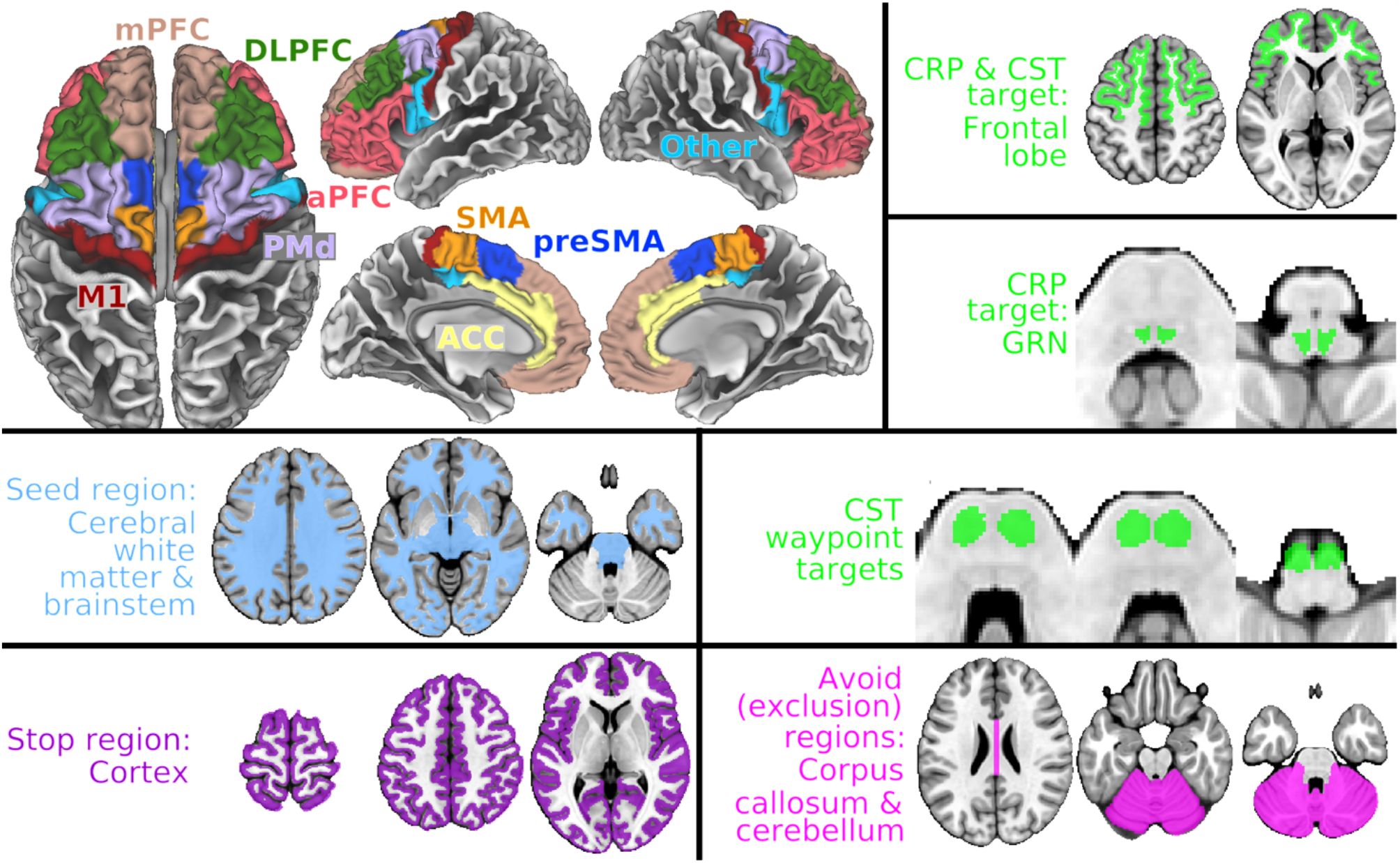
Tractography regions. <u>Top left panel</u>: Frontal cortex streamline inclusion region on the gray/white matter boundary surface, with subregions labelled by color. This region was mapped to volume space and dilated 2 mm inward for tractography (top right panel), as were its subregions. <u>Remaining panels</u>: streamline inclusion regions (lime green), seeding region (pale blue), stopping region (purple) and exclusion regions (fuscia), in the MNI tractography space (MNI152NLin2009cAsym). ACC; anterior cingulate cortex; aPFC, anterior prefrontal cortex; CRP, corticoreticular pathway; CST, corticospinal tract; DLPFC, dorsolateral prefrontal cortex; GRN, gigantocellular reticular nucleus; M1, primary motor cortex; mPFC, medial prefrontal cortex; PMd, dorsal premotor cortex; SMA, supplementary motor area. Data at: https://balsa.wustl.edu/L65ZX (upon acceptance for publication).

### 3.2. CRP and CST diffusion tractography results (Figure 3)

#### 3.2.1. Cortical endpoints

CRP streamlines were distributed throughout widespread parts of both frontal lobes, with major cortical endpoints in M1, SMA, preSMA, PMd, mPFC and aPFC regions. CST streamlines had cortical endpoints primarily in the M1, SMA, preSMA and PMd regions. The CRP and CST streamline endpoints partially overlapped in most subregions of the motor cortex. Compared with the CST, CRP streamlines were more distributed throughout the frontal lobe, sparser for M1, similar for PMd & SMA, and denser for more anterior regions.

**Figure 3.**
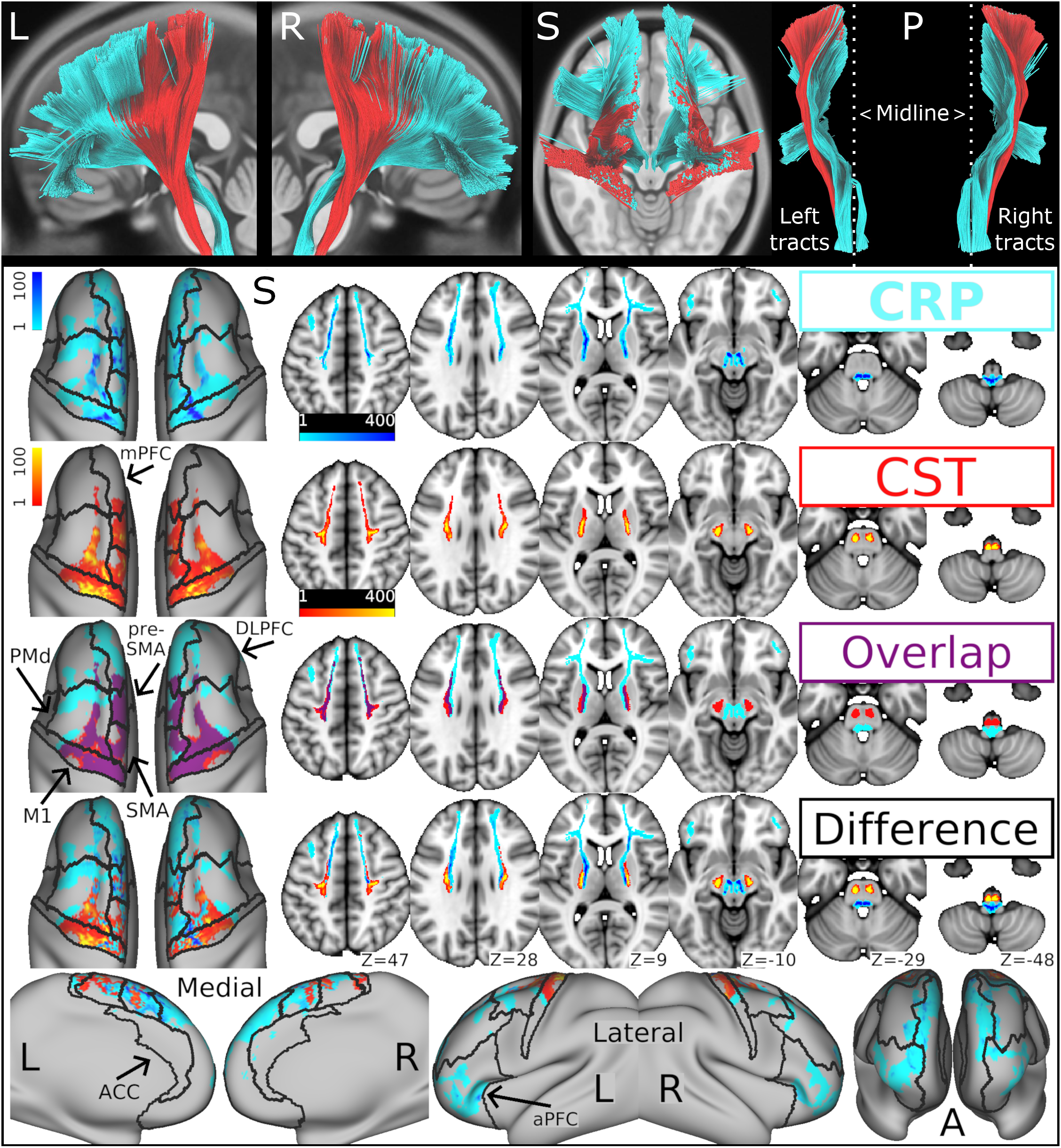
Human average corticoreticular pathway (CRP) template in comparison with the corticospinal tract (CST). The CRP is shown in blue and the CST in red. <u>Upper panel</u> (black background): 3-dimensional rendering of the diffusion streamlines from left (L), right (R), superior (S) and posterior (P) views, in the MNI tractography space (MNI152NLin2009cAsym). In the posterior view, the left and right tracts are separated, and the sagittal midline is marked with a dotted line to visualize contralateral CRP projections. <u>Lower panel</u> (white background): The CRP and CST rows are streamline density images for the respective tracts. The ‘Overlap’ row shows each tract binarized with their overlap area in purple. The bottom two ‘Difference’ rows show subtraction images with the difference in streamline density between the CRP and CST. The streamline density data are shown projected onto an inflated surface model of the cerebral cortex (left column & bottom row) and in the FSL MNI152 template volume (remaining images). Black outlines on the surface models show the frontal lobe subregions. The upper four rows show images from a superior (S) view. The bottom row shows images from medial, lateral and anterior (A) views. ACC; anterior cingulate cortex; aPFC, anterior prefrontal cortex; DLPFC, dorsolateral prefrontal cortex; M1, primary motor cortex; mPFC, medial prefrontal cortex; PMd, dorsal premotor cortex; SMA, supplementary motor area. Data at: https://balsa.wustl.edu/pkj79 (upon acceptance for publication).

#### 3.2.2. Subcortical trajectories

In the subcortex, CRP streamlines were found in the corona radiata (anterior & superior portions), anterior & posterior limbs of the internal capsule (ALIC & PLIC), medioventral diencephalon, midbrain tegmentum and posterior reticular formation regions of the pons and medulla. CST streamlines were found in the superior corona radiata, PLIC, cerebral peduncles and pyramids. The CRP and CST streamlines partially overlapped in the superior corona radiata and PLIC. Compared with the CST, the CRP streamlines were more distributed in the cerebral white matter, more anterior & medial in the internal capsule, more medial in the ventral diencephalon and more posterior in the brainstem.

When filtering CRP streamlines to only include those that touched the left frontal lobe target, some streamlines reached the left GRN target and others reached the right GRN target, with decussation in the midbrain tegmentum (Fig 3, top right panel). The same was true when filtering CRP streamlines to only include those that touched the right frontal lobe target.

## 4. DISCUSSION

This study aimed to map the normal human CRP, using multimodal delineation of the primary reticulospinal nucleus target (GRN) and high-resolution diffusion tractography with group-average brain MRI data (N=1,065), while also including CST tractography for comparison. We were able to manually label the GRN at an adequate level of confidence, with most of its borders visible on at least one MRI contrast. We were also successful in reconstructing a CRP and CST estimate from each hemisphere. The estimated CRP originated from widespread cortical regions, with dense projections from M1, SMA, preSMA, PMd, mPFC and aPFC, while the estimated CST mainly originated from M1, SMA, preSMA and PMd, as expected.[8,77,78] The estimated CST followed its known trajectory through the superior corona radiata, PLIC and cerebral peduncles.[8,78–80] Meanwhile, the estimated CRP had a more distributed and anteromedial cerebral trajectory through both the anterior & superior portions of the corona radiata and through both the ALIC & PLIC, before moving posterior to the CST in the midbrain tegmentum and converging on the pontomedullary reticular formation.

Our results provide the first human average templates of the GRN and CRP in standard MRI space and will be made publicly available at https://balsa.wustl.edu/study/show/v83MM upon acceptance of this manuscript for publication. The CRP templates may be particularly useful for CRP assessment in clinical populations, for whom diffusion tractography is not always feasible or reliable (e.g. failed propagation through an area of pathology may result in streamlines not meeting acceptance criteria even for intact pathways).[79,81] With accurate registration between individual and standard MRI space, a group-level tract template can be aligned with an individual MRI to validly extract data (e.g. anisotropy values, lesion overlap) from the tract of interest using automated software pipelines, even in cases of substantial pathology.[79,81]

Group-level maps of the normal CST are already commonly used in this way.^e.g.^[21,81–86] For example, one recent stroke study used this method to assess the CST, but were forced to do individual diffusion tractography for the CRP due to the lack of an available CRP template.[21] The resultant tractography maps from the lesioned hemisphere were not shown, and CRP streamlines from the non-lesioned hemisphere did not appear to reach the cortex in the image provided.[21] Our tract templates could facilitate this type of research, enabling advantageous CST assessment methods to be similarly applied to the CRP and avoiding the need for individual tractography. We provide full CRP and CST tract density images without binarizing, so that they may be used as weighting maps when extracting averaged data from each tract.[81,86] We also provide versions of each tract template that are restricted to the internal capsule (e.g. CRP_L_IC.nii.gz), since data extraction from this region of the CST is common,^e.g.^[21,81–83] better differentiates patients with stroke from controls,[81] and has been found to be more strongly associated with motor function.[81]

One novel and important finding in the current study was the presence of human CRP streamlines appearing to originate far anterior to the cortical motor areas, from distributed regions of the medial and anterior prefrontal cortices. This finding is reinforced by a recent axonal tracing analysis in the mouse that also identified dense CRP projections from the medial prefrontal cortex (including some analogues of the human anterior prefrontal cortex).[7] That axonal tracing study additionally found dense CRP projections from the anterior cingulate cortex,[7] which were not clearly identified in the current CRP map (although streamlines were present near the anterior cingulate region). One possible explanation for this apparent discrepancy is that any such projections would have likely been passing through the dense cingulum bundle at a relatively sharp turning angle. If so, they would be particularly difficult to track with diffusion tractography.[70]

Regardless, the presence of CRP projections originating far anterior to the motor cortex and passing through the anterior corona radiata & ALIC theoretically increases the likelihood of at least partial CRP sparing after brain lesions and may provide novel targets for neuromodulation. These projections could also explain previously observed associations between medial prefrontal cortex activity and motor function.[40–45] Future studies to better understand the potential motor relevance of such anterior CRP projections should be a priority.

The current CRP map also included novel streamlines appearing to originate from bilateral regions of the anterolateral prefrontal cortex, partially overlapping with the anterior frontal operculum, anterior inferior frontal sulcus and Brodmann areas 45, 47, 9-46v and 10 (Fig 3, bottom row, lateral views). This projection has not been observed in prior animal axonal tracing studies,[7] possibly because the anterior prefrontal cortex is uniquely well developed in humans.[87] However, without any anatomical validation, it is unclear whether these streamlines could represent novel CRP projections (e.g. to central pattern generators for orofacial, autonomic or somatic motor functions[88]) or false pathways.

Another important finding was that the estimated CRP and CST origins overlapped across most of the motor cortex, except the anterior PMd (Fig 3). Primary motor cortex stimulation or recording is often assumed to specifically target the CST.^e.g.^[89–92] Yet, the current results suggest that such activities target both the CST and CRP. This is strongly supported by invasive brainstem recording during cortical stimulation in primates,[39] and it has implications for interpreting the results of many neurophysiologic studies. The current findings also suggest that there is no substantial cortical region that is specific to the CST, while more anterior cortical regions may be more specific to the CRP (vs. the CST). Future neurophysiologic studies are needed to test this hypothesis. Interestingly, our results indicate that even the hand region of the primary motor cortex contributes CRP projections. This conflicts with the persistent traditional view of the CReST as a pathway serving only trunk and proximal limb muscles.[35] However, it is consistent with a large body of evidence in humans and other primates indicating that the CRP is also involved with distal limb function.[93]

This study also provided the first assessment of human CRP laterality. The results appear to confirm that both the left and right CRP project bilaterally to the brainstem, as consistently observed in invasive animal studies across species (including primates).[5–7,36,46,47] The estimated human CRP decussation point in the midbrain tegmentum was more focal than prior decussation estimates in the mouse, which spanned most of the brainstem.[7] It is unclear whether this apparent discrepancy is due to differences in methodology or evolutionary changes. Nevertheless, the existence of these bilateral CRP projections in humans has important implications for recovery from unilateral supratentorial brain damage (e.g. typical stroke). For example, the effects of contralesional hemisphere upregulation after stroke could be at least partially mediated through the contralateral CRP projection from the contralesional cortex to the ipsilesional brainstem.[2,7,24]

### 4.1. Strengths and Limitations

Compared with prior human CRP tractography studies, this work features several methodologic advances to improve accuracy & reliability, including: greater sample size, upgraded MRI hardware,[51,94] higher spatial & angular resolution,[51,74,94] better distortion correction,[51,53,74] advanced model-free reconstruction of diffusion orientations (for improved resolution of multiple fiber populations within a voxel and partial volume effects),[30,55,72,95–97] group-averaging of diffusion orientations prior to tractography,[30,54] and more anatomically-informed delineation of tractography regions[2,7,27–29,48,50,74] (Table 1).

However, even with these careful state-of-the-art methods, diffusion tractography is still an imperfect method for tract tracing, because of insufficient resolution to resolve the microscopic anatomy and inability to differentiate afferent from efferent fibers.[74] Thus, it is possible that our CRP map includes some false streamlines and may still be missing some parts of the true CRP. For example, diffusion tractography is known to have more difficulty resolving fibers projecting from sulci vs. gyri, on average.[69] A likely example of this can be seen in the primary motor cortex hand knob near the central sulcus, where no CRP or CST streamlines were found (Figure 3: left column, middle rows), even though this is a known origin of at least CST axons.

Another limitation is that we did not attempt to map CRP or CST projections from the somatosensory cortex, which have been identified in invasive animal studies.[7] This was done to minimize false streamlines tracing sensory pathways in the tract maps, and because the somatosensory cortex did not make a primary contribution to either tract in previous research.[7] We also could not fully assess for the presence of CRP projections arising as collateral branches of CST axons as observed in animal studies,[3,4,36,38] because the tractography SDF map did not include enough of the medulla. If such CRP projections are present in humans, our current methods would have underestimated the density of CRP projections from the motor cortex. Lastly, the diffusion MRI data used for tractography were exclusively from young adults, so future studies may wish to test for age-related changes to the CRP origins and trajectory. However, CST location appears to be stable during aging[98] and we have no reason to believe that CRP location would be any less stable.

### 4.2. Conclusions

The human CRP appears to originate from large portions of the frontal cortex, including M1, SMA, preSMA, PMd, mPFC and aPFC, with distributed projections through the anterior & superior corona radiata and the ALIC & PLIC before partially decussating in the midbrain tegmentum and converging bilaterally on the pontomedullary reticular formation. CRP and CST origins seem to overlap for most of the motor cortex, while the CRP appears to have inputs from more anterior regions of the frontal lobe that are not present for the CST. The CRP also seems to have a more distributed cerebral trajectory, which was partially overlapping and partially anteromedial to the CST. These findings have important implications for motor recovery after brain lesions. In addition, the GRN and CRP templates generated from this study should facilitate future research in this area.

## ABBREVIATIONS

ALIC: anterior limb of internal capsule
aPFC: anterior prefrontal cortex
CReST: cortico-reticulo-spinal tract
CRP: corticoreticular pathway
CST: corticospinal tract
DEC: direction-encoded color
DLPFC: dorsolateral prefrontal cortex
EPI: echo-planar imaging
FA: fractional anisotropy
FSL: FMRIB software library
GRN: gigantocellular reticular nucleus
HCP: human connectome project
M1: primary motor cortex
MNI: Montreal Neurologic Institute
mPFC: medial prefrontal cortex
PLIC: posterior limb of internal capsule
PMd: dorsal premotor cortex
PMv: ventral premotor cortex
QA: quantitative anisotropy
SDF: spin distribution function
SMA: supplementary motor area
RST: reticulospinal tract

## REFERENCES

[1] Shapovalov AI. Evolution of neuronal systems of suprasegmental motor control. Neurophysiology (New York) 1973;4:346–59.

[2] Brownstone RM, Chopek JW. Reticulospinal systems for tuning motor commands. Frontiers in neural circuits 2018;12:30.

[3] Jinnai K. Electrophysiological study on the corticoreticular projection neurons of the cat. Brain Res 1984;291:145–9.

[4] Matsuyama K, Mori F, Nakajima K, Drew T, Aoki M, Mori S. Locomotor role of the corticoreticular-reticulospinal-spinal interneuronal system. Prog Brain Res 2004;143:239–49.

[5] Fisher KM, Zaaimi B, Edgley SA, Baker SN. Extensive Cortical Convergence to Primate Reticulospinal Pathways. J Neurosci 2021;41:1005–18.

[6] Fregosi M, Contestabile A, Hamadjida A, Rouiller EM. Corticobulbar projections from distinct motor cortical areas to the reticular formation in macaque monkeys. Eur J Neurosci 2017;45:1379–95.

[7] Boyne P, Awosika OO, Luo Y. Mapping the corticoreticular pathway from cortex-wide anterograde axonal tracing in the mouse. bioRxiv 449661 [Preprint] 2021.

[8] Welniarz Q, Dusart I, Roze E. The corticospinal tract: Evolution, development, and human disorders. Developmental neurobiology (Hoboken, N.J.) 2017;77:810–29.

[9] Castle-Kirszbaum M, Goldschlager T. Pyramidal weakness: Is it time to retire the term? Clin Anat 2021;34:478–82.

[10] McPherson JG, Chen A, Ellis MD, Yao J, Heckman CJ, Dewald JPA. Progressive recruitment of contralesional cortico-reticulospinal pathways drives motor impairment post stroke. J Physiol (Lond) 2018;596:1211–25.

[11] Karbasforoushan H, Cohen-Adad J, Dewald JPA. Brainstem and spinal cord MRI identifies altered sensorimotor pathways post-stroke. Nature communications 2019;10:3524.

[12] Ellis MD, Drogos J, Carmona C, Keller T, Dewald JPA. Neck rotation modulates flexion synergy torques, indicating an ipsilateral reticulospinal source for impairment in stroke. J Neurophysiol 2012;108:3096–104.

[13] Zaaimi B, Soteropoulos DS, Fisher KM, Riddle CN, Baker SN. Classification of Neurons in the Primate Reticular Formation and Changes after Recovery from Pyramidal Tract Lesion. J Neurosci 2018;38:6190–206.

[14] Asboth L, Friedli L, Beauparlant J, Martinez-Gonzalez C, Anil S, Rey E, Baud L, Pidpruzhnykova G, Anderson MA, Shkorbatova P, Batti L, Pages S, Kreider J, Schneider BL, Barraud Q, Courtine G. Cortico-reticulo-spinal circuit reorganization enables functional recovery after severe spinal cord contusion. Nat Neurosci 2018;21:576–88.

[15] Zaaimi B, Edgley SA, Soteropoulos DS, Baker SN. Changes in descending motor pathway connectivity after corticospinal tract lesion in macaque monkey. Brain 2012;135:2277–89.

[16] Glover IS, Baker SN. Cortical, Corticospinal, and Reticulospinal Contributions to Strength Training. J Neurosci 2020;40:5820–32.

[17] Takase H, Kurihara Y, Yokoyama TA, Kawahara N, Takei K. LOTUS overexpression accelerates neuronal plasticity after focal brain ischemia in mice. PLoS One 2017;12:e0184258.

[18] Darling WG, Ge J, Stilwell-Morecraft KS, Rotella DL, Pizzimenti MA, Morecraft RJ. Hand motor recovery following extensive frontoparietal cortical injury is accompanied by upregulated corticoreticular projections in monkey. The Journal of neuroscience 2018;38:6323–39.

[19] Ishida A, Kobayashi K, Ueda Y, Shimizu T, Tajiri N, Isa T, Hida H. Dynamic Interaction between Cortico-Brainstem Pathways during Training-Induced Recovery in Stroke Model Rats. The Journal of neuroscience 2019;39:7306–20.

[20] Jang SH, Chang CH, Lee J, Kim CS, Seo JP, Yeo SS. Functional role of the corticoreticular pathway in chronic stroke patients. Stroke 2013;44:1099–104.

[21] Soulard J, Huber C, Baillieul S, Thuriot A, Renard F, Aubert Broche B, Krainik A, Vuillerme N, Jaillard A. Motor tract integrity predicts walking recovery. Neurology 2020;94:e583–93.

[22] Yeo SS, Jang SH, Park GY, Oh S. Effects of injuries to descending motor pathways on restoration of gait in patients with pontine hemorrhage. J Stroke Cerebrovasc Dis 2020;29:104857.

[23] Jun S, Hong B, Kim Y, Lim S. Does Motor Tract Integrity at 1 Month Predict Gait and Balance Outcomes at 6 Months in Stroke Patients? Brain Sciences 2021;11.

[24] Cleland BT, Madhavan S. Ipsilateral motor pathways to the lower limb after stroke: Insights and opportunities. J Neurosci Res 2021;99:1565–78.

[25] Taga M, Charalambous CC, Raju S, Lin J, Zhang Y, Stern E, Schambra HM. Corticoreticulospinal tract neurophysiology in an arm and hand muscle in healthy and stroke subjects. J Physiol (Lond) 2021:In press.

[26] Hirabayashi R, Kojima S, Edama M, Onishi H. Activation of the supplementary motor areas enhances spinal reciprocal inhibition in healthy individuals. Brain sciences 2020;10:587.

[27] Azadbakht H, Parkes LM, Haroon HA, Augath M, Logothetis NK, de Crespigny A, D’Arceuil HE, Parker GJ. Validation of High-Resolution Tractography Against In Vivo Tracing in the Macaque Visual Cortex. Cereb Cortex 2015;25:4299–309.

[28] Aydogan DB, Jacobs R, Dulawa S, Thompson SL, Francois MC, Toga AW, Dong H, Knowles JA, Shi Y. When tractography meets tracer injections: a systematic study of trends and variation sources of diffusion-based connectivity. Brain Struct Funct 2018;223:2841–58.

[29] Gutierrez CE, Skibbe H, Nakae K, Tsukada H, Lienard J, Watakabe A, Hata J, Reisert M, Woodward A, Yamaguchi Y, Yamamori T, Okano H, Ishii S, Doya K. Optimization and validation of diffusion MRI-based fiber tracking with neural tracer data as a reference. Sci Rep 2020;10:21285–4.

[30] Yeh F, Panesar S, Fernandes D, Meola A, Yoshino M, Fernandez-Miranda JC, Vettel JM, Verstynen T. Population-averaged atlas of the macroscale human structural connectome and its network topology. NeuroImage (Orlando, Fla.) 2018;178:57–68.

[31] Yeo SS, Chang MC, Kwon YH, Jung YJ, Jang SH. Corticoreticular pathway in the human brain: diffusion tensor tractography study. Neurosci Lett 2012;508:9–12.

[32] Yeo SS, Jang SH, Son SM. The different maturation of the corticospinal tract and corticoreticular pathway in normal brain development: diffusion tensor imaging study. Front Hum Neurosci 2014;8:573.

[33] Jang SH, Seo JP. The distribution of the cortical origin of the corticoreticular pathway in the human brain: a diffusion tensor imaging study. Somatosens Mot Res 2014;31:204–8.

[34] Jang SH, Seo JP. The anatomical location of the corticoreticular pathway at the subcortical white matter in the human brain: A diffusion tensor imaging study. Somatosens Mot Res 2015;32:106–9.

[35] Jang SH, Lee SJ. Corticoreticular Tract in the Human Brain: A Mini Review. Front Neurol 2019;10:1188.

[36] Kably B, Drew T. Corticoreticular pathways in the cat. I. Projection patterns and collaterization. J Neurophysiol 1998;80:389–405.

[37] He X, Wu C. Connections between pericruciate cortex and the medullary reticulospinal neurons in cat: an electrophysiological study. Experimental brain research 1985;61:109–16.

[38] Lamas JA, Martinez L, Canedo A. Pericruciate fibres to the red nucleus and to the medial bulbar reticular formation. Neuroscience 1994;62:115–24.

[39] Fisher KM, Zaaimi B, Baker SN. Reticular formation responses to magnetic brain stimulation of primary motor cortex. J Physiol 2012;590:4045–60.

[40] Boyne P, Maloney T, DiFrancesco M, Fox MD, Awosika O, Aggarwal P, Woeste J, Jaroch L, Braswell D, Vannest J. Resting-state functional connectivity of subcortical locomotor centers explains variance in walking capacity. Hum Brain Mapp 2018;39:4831–43.

[41] Gwin JT, Gramann K, Makeig S, Ferris DP. Electrocortical activity is coupled to gait cycle phase during treadmill walking. Neuroimage 2011;54:1289–96.

[42] Knaepen K, Mierau A, Tellez HF, Lefeber D, Meeusen R. Temporal and spatial organization of gait-related electrocortical potentials. Neurosci Lett 2015;599:75–80.

[43] Tian Q, Chastan N, Bair WN, Resnick SM, Ferrucci L, Studenski SA. The brain map of gait variability in aging, cognitive impairment and dementia-A systematic review. Neurosci Biobehav Rev 2017;74:149–62.

[44] Boyne P, Doren S, Scholl V, Staggs E, Whitesel D, Maloney T, Awosika O, Kissela B, Dunning K, Vannest J. Functional magnetic resonance brain imaging of imagined walking to study locomotor function after stroke. Clin Neurophysiol 2020;132:167–77.

[45] Cremers J, D’Ostilio K, Stamatakis J, Delvaux V, Garraux G. Brain activation pattern related to gait disturbances in Parkinson’s disease. Mov Disord 2012;27:1498–505.

[46] Matsuyama K, Drew T. Organization of the projections from the pericruciate cortex to the pontomedullary brainstem of the cat: a study using the anterograde tracer Phaseolus vulgaris-leucoagglutinin. J Comp Neurol 1997;389:617–41.

[47] Rho M, Cabana T, Drew T. Organization of the projections from the pericruciate cortex to the pontomedullary reticular formation of the cat: A quantitative retrograde tracing study. Journal of comparative neurology (1911) 1997;388:228–49.

[48] Lechanoine F, Jacquesson T, Beaujoin J, Serres B, Mohammadi M, Planty-Bonjour A, Andersson F, Poupon F, Poupon C, Destrieux C. WIKIBrainStem: An online atlas to manually segment the human brainstem at the mesoscopic scale from ultrahigh field MRI. NeuroImage (Orlando, Fla.) 2021;236:118080.

[49] Van Essen DC, Smith SM, Barch DM, Behrens TE, Yacoub E, Ugurbil K, WU-Minn HCP Consortium. The WU-Minn Human Connectome Project: an overview. Neuroimage 2013;80:62–79.

[50] Paxinos G, Xu-Feng H, Sengul G, Watson C. Organization of Brainstem Nuclei. 2012:260–327.

[51] Sotiropoulos SN, Jbabdi S, Xu J, Andersson JL, Moeller S, Auerbach EJ, Glasser MF, Hernandez M, Sapiro G, Jenkinson M, Feinberg DA, Yacoub E, Lenglet C, Van Essen DC, Ugurbil K, Behrens TE, WU-Minn HCP Consortium. Advances in diffusion MRI acquisition and processing in the Human Connectome Project. Neuroimage 2013;80:125–43.

[52] Van Essen DC, Ugurbil K, Auerbach E, Barch D, Behrens TE, Bucholz R, Chang A, Chen L, Corbetta M, Curtiss SW, Della Penna S, Feinberg D, Glasser MF, Harel N, Heath AC, Larson-Prior L, Marcus D, Michalareas G, Moeller S, Oostenveld R, Petersen SE, Prior F, Schlaggar BL, Smith SM, Snyder AZ, Xu J, Yacoub E, WU-Minn HCP Consortium. The Human Connectome Project: a data acquisition perspective. Neuroimage 2012;62:2222–31.

[53] Glasser MF, Sotiropoulos SN, Wilson JA, Coalson TS, Fischl B, Andersson JL, Xu J, Jbabdi S, Webster M, Polimeni JR, Van Essen DC, Jenkinson M, WU-Minn HCP Consortium. The minimal preprocessing pipelines for the Human Connectome Project. Neuroimage 2013;80:105–24.

[54] Yeh F, Tseng WI. NTU-90: A high angular resolution brain atlas constructed by q-space diffeomorphic reconstruction. NeuroImage (Orlando, Fla.) 2011;58:91–9.

[55] Yeh F, Verstynen TD, Wang Y, Fernández-Miranda JC, Tseng WI. Deterministic Diffusion Fiber Tracking Improved by Quantitative Anisotropy. PloS one 2013;8:e80713.

[56] Liang H, Paxinos G, Watson C. Projections from the brain to the spinal cord in the mouse. Brain structure & function 2011;215:159–86.

[57] Torvik A, Brodal A. The origin of reticulospinal fibers in the cat. An experimental study. Anat Rec 1957;128:113–37.

[58] Horn AKE, Adamczyk C. Reticular Formation : Eye Movements, Gaze and Blinks. 2012:328–66.

[59] Jenkinson M, Beckmann CF, Behrens TE, Woolrich MW, Smith SM. Fsl. Neuroimage 2012;62:782–90.

[60] Jenkinson M, Bannister P, Brady M, Smith S. Improved Optimization for the Robust and Accurate Linear Registration and Motion Correction of Brain Images. NeuroImage (Orlando, Fla.) 2002;17:825–41.

[61] Andersson J, Jenkinson M, Smith S. Non-linear registration a.k.a. spatial normalisation. FMRIB Technial Report TR07JA2. 2007.

[62] Fischl B. FreeSurfer. Neuroimage 2012;62:774–81.

[63] Desikan RS, Ségonne F, Fischl B, Quinn BT, Dickerson BC, Blacker D, Buckner RL, Dale AM, Maguire RP, Hyman BT, Albert MS, Killiany RJ. An automated labeling system for subdividing the human cerebral cortex on MRI scans into gyral based regions of interest. NeuroImage (Orlando, Fla.) 2006;31:968–80.

[64] Klein A, Tourville J. 101 labeled brain images and a consistent human cortical labeling protocol. Frontiers in neuroscience 2012;6:171.

[65] Glasser MF, Coalson TS, Robinson EC, Hacker CD, Harwell J, Yacoub E, Ugurbil K, Andersson J, Beckmann CF, Jenkinson M, Smith SM, Van Essen DC. A multi-modal parcellation of human cerebral cortex. Nature 2016;536:171–8.

[66] Van Essen DC, Glasser MF, Dierker DL, Harwell J, Coalson T. Parcellations and hemispheric asymmetries of human cerebral cortex analyzed on surface-based atlases. Cereb Cortex 2012;22:2241–62.

[67] Robinson EC, Garcia K, Glasser MF, Chen Z, Coalson TS, Makropoulos A, Bozek J, Wright R, Schuh A, Webster M, Hutter J, Price A, Cordero Grande L, Hughes E, Tusor N, Bayly PV, Van Essen DC, Smith SM, Edwards AD, Hajnal J, Jenkinson M, Glocker B, Rueckert D. Multimodal surface matching with higher-order smoothness constraints. Neuroimage 2018;167:453–65.

[68] Schilling KG, Gao Y, Stepniewska I, Janve V, Landman BA, Anderson AW. Anatomical accuracy of standard-practice tractography algorithms in the motor system - A histological validation in the squirrel monkey brain. Magn Reson Imaging 2019;55:7–25.

[69] Schilling K, Gao Y, Janve V, Stepniewska I, Landman BA, Anderson AW. Confirmation of a gyral bias in diffusion MRI fiber tractography. Hum Brain Mapp 2018;39:1449–66.

[70] Reveley C, Seth AK, Pierpaoli C, Silva AC, Yu D, Saunders RC, Leopold DA, Ye FQ. Superficial white matter fiber systems impede detection of long-range cortical connections in diffusion MR tractography. Proceedings of the National Academy of Sciences - PNAS 2015;112:E2820–8.

[71] Fischl B, Salat DH, Busa E, Albert M, Dieterich M, Haselgrove C, van der Kouwe A, Killiany R, Kennedy D, Klaveness S, Montillo A, Makris N, Rosen B, Dale AM. Whole Brain Segmentation: Automated Labeling of Neuroanatomical Structures in the Human Brain. Neuron (Cambridge, Mass.) 2002;33:341–55.

[72] Maier-Hein KH, Neher PF, Houde J, Côté M, Garyfallidis E, Zhong J, Chamberland M, Yeh F, Lin Y, Ji Q, Reddick WE, Glass JO, Chen DQ, Feng Y, Gao C, Wu Y, Ma J, Renjie H, Li Q, Westin C, Deslauriers-Gauthier S, González JOO, Paquette M, St-Jean S, Girard G, Rheault F, Sidhu J, Tax CMW, Guo F, Mesri HY, Dávid S, Froeling M, Heemskerk AM, Leemans A, Boré A, Pinsard B, Bedetti C, Desrosiers M, Brambati S, Doyon J, Sarica A, Vasta R, Cerasa A, Quattrone A, Yeatman J, Khan AR, Hodges W, Alexander S, Romascano D, Barakovic M, Auría A, Esteban O, Lemkaddem A, Thiran J, Cetingul HE, Odry BL, Mailhe B, Nadar MS, Pizzagalli F, Prasad G, Villalon-Reina JE, Galvis J, Thompson PM, Requejo FDS, Laguna PL, Lacerda LM, Barrett R, Dell’Acqua F, Catani M, Petit L, Caruyer E, Daducci A, Dyrby TB, Holland-Letz T, Hilgetag CC, Stieltjes B, Descoteaux M. The challenge of mapping the human connectome based on diffusion tractography. Nature communications 2017;8:1349–13.

[73] Yeh FC. Shape analysis of the human association pathways. Neuroimage 2020;223:117329.

[74] Calamante F. The Seven Deadly Sins of Measuring Brain Structural Connectivity Using Diffusion MRI Streamlines Fibre-Tracking. Diagnostics (Basel) 2019;9:115.

[75] Nathan PW, Smith MC, Deacon P. The corticospinal tracts in man. Course and location of fibres at different segmental levels. Brain (London, England : 1878) 1990;113:303–24.

[76] Mayka MA, Corcos DM, Leurgans SE, Vaillancourt DE. Three-dimensional locations and boundaries of motor and premotor cortices as defined by functional brain imaging: A meta-analysis. NeuroImage (Orlando, Fla.) 2006;31:1453–74.

[77] Seo JP, Jang SH. Different characteristics of the corticospinal tract according to the cerebral origin: DTI study. American journal of neuroradiology : AJNR 2013;34:1359–63.

[78] Archer DB, Vaillancourt DE, Coombes SA. A Template and Probabilistic Atlas of the Human Sensorimotor Tracts using Diffusion MRI. Cereb Cortex 2018;28:1685–99.

[79] Hua K, Zhang J, Wakana S, Jiang H, Li X, Reich DS, Calabresi PA, Pekar JJ, van Zijl, Peter C. M., Mori S. Tract probability maps in stereotaxic spaces: Analyses of white matter anatomy and tract-specific quantification. NeuroImage (Orlando, Fla.) 2008;39:336–47.

[80] Wakana S, Caprihan A, Panzenboeck MM, Fallon JH, Perry M, Gollub RL, Hua K, Zhang J, Jiang H, Dubey P, Blitz A, van Zijl P, Mori S. Reproducibility of quantitative tractography methods applied to cerebral white matter. NeuroImage (Orlando, Fla.) 2007;36:630–44.

[81] Park CH, Kou N, Boudrias MH, Playford ED, Ward NS. Assessing a standardised approach to measuring corticospinal integrity after stroke with DTI. Neuroimage Clin 2013;2:521–33.

[82] Stinear CM, Barber PA, Smale PR, Coxon JP, Fleming MK, Byblow WD. Functional potential in chronic stroke patients depends on corticospinal tract integrity. Brain 2007;130:170–80.

[83] Stinear CM, Barber PA, Petoe M, Anwar S, Byblow WD. The PREP algorithm predicts potential for upper limb recovery after stroke. Brain (London, England : 1878) 2012;135:2527–35.

[84] Dawes H, Enzinger C, Johansen-Berg H, Bogdanovic M, Guy C, Collett J, Izadi H, Stagg C, Wade D, Matthews PM. Walking performance and its recovery in chronic stroke in relation to extent of lesion overlap with the descending motor tract. Exp Brain Res 2008;186:325–33.

[85] Schulz R, Park C, Boudrias M, Gerloff C, Hummel FC, Ward NS. Assessing the integrity of corticospinal pathways from primary and secondary cortical motor areas after stroke. Stroke (1970) 2012;43:2248–51.

[86] Riley JD, Le V, Der-Yeghiaian L, See J, Newton JM, Ward NS, Cramer SC. Anatomy of stroke injury predicts gains from therapy. Stroke 2011;42:421–6.

[87] Carlen M. What constitutes the prefrontal cortex? Science (American Association for the Advancement of Science) 2017;358:478–82.

[88] Steuer I, Guertin PA. Central pattern generators in the brainstem and spinal cord: An overview of basic principles, similarities and differences. Rev Neurosci 2019;30:107–64.

[89] Barthélemy D, Grey MJ, Nielsen JB, Bouyer L. Involvement of the corticospinal tract in the control of human gait. Prog Brain Res 2011;192:181.

[90] Jayaram G, Stagg CJ, Esser P, Kischka U, Stinear J, Johansen-Berg H. Relationships between functional and structural corticospinal tract integrity and walking post stroke. Clin Neurophysiol 2012;123:2422–8.

[91] Capaday C, Lavoie BA, Barbeau H, Schneider C, Bonnard M. Studies on the Corticospinal Control of Human Walking. I. Responses to Focal Transcranial Magnetic Stimulation of the Motor Cortex. J Neurophysiol 1999;81:129–39.

[92] Chieffo R, Comi G, Leocani L. Noninvasive Neuromodulation in Poststroke Gait Disorders: Rationale, Feasibility, and State of the Art. Neurorehabilitation and Neural Repair 2016;30:71–82.

[93] Baker SN, Zaaimi B, Fisher KM, Edgley SA, Soteropoulos DS. Pathways mediating functional recovery. Prog Brain Res 2015;218:389–412.

[94] Ugurbil K, Xu J, Auerbach EJ, Moeller S, Vu AT, Duarte-Carvajalino JM, Lenglet C, Wu X, Schmitter S, Van de Moortele, p F, Strupp J, Sapiro G, De Martino F, Wang D, Harel N, Garwood M, Chen L, Feinberg DA, Smith SM, Miller KL, Sotiropoulos SN, Jbabdi S, Andersson JL, Behrens TE, Glasser MF, Van Essen DC, Yacoub E, WU-Minn HCP Consortium. Pushing spatial and temporal resolution for functional and diffusion MRI in the Human Connectome Project. Neuroimage 2013;80:80–104.

[95] Yeh F, Wedeen VJ, Tseng WI. Generalized q-sampling imaging. IEEE Trans Med Imaging 2010;29:1626–35.

[96] Yeh F, Tseng WI. Sparse Solution of Fiber Orientation Distribution Function by Diffusion Decomposition. PloS one 2013;8:e75747.

[97] Gangolli M, Holleran L, Hee Kim J, Stein TD, Alvarez V, McKee AC, Brody DL. Quantitative validation of a nonlinear histology-MRI coregistration method using generalized Q-sampling imaging in complex human cortical white matter. NeuroImage (Orlando, Fla.) 2017;153:152–67.

[98] Jang SH, Seo JP. Aging of corticospinal tract fibers according to the cerebral origin in the human brain: A diffusion tensor imaging study. Neurosci Lett 2015;585:77–81.

